# Comparing the efficacy of data-driven denoising methods for a multi-echo fMRI acquisition at 7T

**DOI:** 10.1101/2022.07.03.498599

**Authors:** Abraham B. Beckers, Benedikt A. Poser, Daniel Keszthelyi

## Abstract

In functional magnetic resonance imaging (fMRI) of the brain the signal is dominated by (physiological) noise. Imaging at ultrahigh field strength is becoming increasingly popular as it offers increased spatial accuracy. The latter is of particular benefit in brainstem neuroimaging given the small cross-sectional area of most nuclei. Physiological noise scales with field strength in fMRI acquisitions, however. Although this problem is in part solved by decreasing voxel size, it is clear that adequate physiological denoising is of utmost importance in brainstem-focused fMRI experiments. Multi-echo sequences have been reported to facilitate highly effective denoising through TE-dependence of Blood Oxygen Level Dependent (BOLD) signals, in a denoising method referred to as multi-echo independent component analysis (ME-ICA). It has not been explored previously how ME-ICA compares to other data-driven denoising approaches at ultrahigh field strength. In the current study, we compared the efficacy of several denoising methods, including anatomical component based correction (aCompCor), Automatic Removal of Motion Artifacts (ICA-AROMA), ME-ICA, and a combination of ME-ICA and aCompCor. We assessed several data quality metrics, including temporal signal-to-noise ratio (tSNR), delta variation signal (DVARS) and spectral density of the global signal. Moreover, we looked at the ability of each method to uncouple the global signal and respiration. In line with previous reports at lower field strengths, we demonstrate that after applying ME-ICA, the data is best post-processed in order to remove spatially diffuse noise with a method such as aCompCor. Our findings indicate that ME-ICA combined with aCompCor and ICA-AROMA are highly effective denoising approaches for multi-echo data acquired at 7T. ME-ICA combined with aCompCor potentially preserves more signal-of-interest as compared to ICA-AROMA.

**Highlights:**

- ME-ICA and ICA-AROMA provide effective denoising for multi-echo 7T fMRI data
- High tSNR can be achieved in the brainstem with a multi-echo acquisition at 7T
- After ME-ICA, the data is best post-processed to correct for spatially diffuse noise

## Introduction

Functional magnetic resonance imaging (fMRI) exploits the small magnetic field distortions induced by deoxyhemoglobin to capture changes in cerebral oxygen supply and, indirectly, neuronal activity (Kwong et al., 1992). To facilitate measurement of these blood oxygenation level-dependent (BOLD) signal changes, fMRI relies on imaging techniques sensitive to susceptibility effects, such as gradient-echo echo-planar imaging (EPI). Given that the fractional change in the MRI signal as a result of most functional tasks does not exceed a few percent, the majority of the temporal signal variation can be classified as noise. The validity of an fMRI experiment that relies on the detection of small task-induced BOLD changes is therefore largely dependent on the efficiency of the applied data cleaning method(s). Various sources of noise can been identified, each warranting its own approach for mitigating confounding effects (Liu, 2016). Common sources of noise include: 1) background noise, for example thermal noise (originating from the subject, and to a lesser extent scanner hardware) and spurious radiofrequency noise from the environment, 2) physiological noise, originating from cardiac and respiratory activity, and low frequency drifts due to intrinsic fluctuations in blood flow and metabolism, and 3) motion-related noise. Many approaches to denoising have been proposed over the past two decades, including methods that simply regress out respiratory and cardiac cycles or locally fit to those fluctuations (Glover et al., 2000), sample at multiple TE (Speck and Hennig, 1998), are based on ICA or PCA and subsequent filtering or a combination of these. Although all fMRI studies are affected by these types of noise, studies focusing of the brainstem can be particularly challenging (Napadow et al., 2019). As a consequence of the close proximity of the basilar artery, pulse pressure waves cause relatively large displacements in the brainstem as compared to cortical regions. Similarly, the adjacent cerebrospinal fluid filled spaces, which flow correlates with the cardiac cycle (Kong et al., 2012), further increases brainstem motion. Being closer to air-filled cavities, such as the sinuses and the chest, also renders the brainstem prone to susceptibility artefacts. The complexity of these artefacts increases due to respiration, as it results in cyclic variability in air content in the chest (Brooks et al., 2013).

Given the small size of most brainstem nuclei – often only a few millimeters wide – high spatial resolutions and tissue specificity are a prerequisite for (functional) brainstem imaging. Scanning at ultra-high field strength (7T and above) can facilitate such high resolutions. One should realize though that – although mitigated by higher spatial resolutions – physiological noise can become an even more dominant source of noise at higher field strengths (Brooks et al., 2013; Triantafyllou et al., 2005). It will be clear that, for a brainstem fMRI experiment to succeed, highly efficient physiological denoising methods are warranted. The performance of such methods at 7T, in particular for brainstem imaging, has not been compared previously.

Previous resting state fMRI studies at 1.5T and 3T have demonstrated that multi-echo EPI (ME-EPI) acquisitions enable the application of more effective data cleaning methods than single echo acquisitions (Dipasquale et al., 2017; Kundu et al., 2012). In addition, acquiring images at multiple echo times reduces signal dropout (Poser and Norris, 2009), which can be particularly problematic in the brainstem. Finally, if an experiment requires scanning both brainstem and cortical areas, the high T2* variability between these areas can be accounted for by the use of T2* maps calculated from ME-EPI data (Puckett et al., 2018).

The aim of the current analysis was to investigate the performance of various data cleaning methods for multiband multi-echo fMRI at 7T in order to increase signal-to-noise ratio, using a dataset recently obtained to investigate brain(stem) responses to an intestinal chemonociceptive stimulus (Beckers et al., 2021). The performance of several data-driven denoising approaches was compared. These methods included anatomical component based correction (aCompCor), Automatic Removal of Motion Artifacts (ICA-AROMA) and multi-echo independent component analysis (ME-ICA). We hypothesized that ME-ICA combined with aCompCor, which fully exploits the benefits of multi-echo imaging, would yield the highest data quality based on the assessed quality metrics.

## Methods

### Participants and data acquisition

This study was approved by the Medical Ethics Committee of the Maastricht University Medical Center/University of Maastricht and registered on clinicaltrials.gov with identifier NCT02551029. Subjects gave written informed consent. Eighteen female healthy volunteers participated in the study (mean age 25 years [±SD 4.4], mean BMI 22.7 kg/m^2^ [±SD 1.9]). Only females were included due to the primary study aims, as described elsewhere in detail (Beckers et al., 2021).

BOLD fMRI data were collected on a Siemens Magnetom 7T scanner (Siemens Healthineers, Erlangen, Germany) using a 32-channel Nova Medical Head Coil. Functional MRI data were acquired with a ME-EPI sequence using an interleaved ascending simultaneous multi-slice acquisition with a multi-band factor of 2. This sequence used the following parameters: 2.2 mm isotropic voxel size, 50 slices, repetition time (TR) = 1.6 s, echo times TE1 = 10.6 ms, TE2 = 26.1 ms, TE3 = 41.6 ms, flip angle = 64°, bandwidth = 2362 Hz/Px, echo spacing = 0.53 ms, GRAPPA acceleration factor R = 3, 1350 volumes with three echoes each. Slices were tilted to fully incorporate the brainstem, the primary focus of the study, while at the same time minimizing the inclusion of air-filled cavities. Because of tilting, image acquisition of part of the parietal cortex was limited up to the postcentral gyrus. Five additional volumes using the same parameters but with opposite phase encoding were acquired for the purpose of susceptibility distortion correction. Concurrent with image acquisition, pulse rate and respiration signals were collected with the use of a pulse oximeter (left index finger) and respiratory belt, respectively.

### Data pre-processing

Data preprocessing was performed using a combination of Statistical Parametric Mapping of the Wellcome Department of Cognitive Neurology of London (SPM12), CONN toolbox (19.b)(Whitfield-Gabrieli and Nieto-Castanon, 2012), the Oxford Centre for Functional MRI of the Brain (FMRIB) Software Library (FSL; v.6.0), and TE Dependent ANAlysis (*tedana* v0.0.8)(Kundu et al., 2012; Kundu et al., 2017).

fMRI images were first corrected for slice timing using manually defined slice timings (extracted from the DICOM header) as applicable for multiband acquisitions, and subsequently motion corrected using an SPM-based script that estimated motion on TE1 images, where after the same transformations were applied to all three echoes. TE-dependence analysis was performed on slice time and motion corrected multi-echo data. An adaptive mask was generated, in which each voxel’s value reflects the number of echoes with usable data. A monoexponential model was fit to the data at each voxel using log-linear regression in order to estimate T2* and S0 maps. For each voxel, the value from the adaptive mask was used to determine which echoes would be used to estimate T2* and S0. Multi-echo data were then optimally combined using the t2s combination method (Posse et al., 1999).

After *tedana*, the preprocessing pipeline branches, providing five different datasets (see Figure 1). The optimally combined non-denoised dataset was corrected for susceptibility-induced distortion (estimated using topup, FSL (Andersson et al., 2003)), segmented and normalized to MNI space. Minimal smoothing using a 4 mm Gaussian kernel was performed as per the preprocessing pipeline described elsewhere (Beckers et al., 2021). The resulting dataset was considered the “uncleaned” dataset, hereafter referred to as ME-uncleaned. The ME-uncleaned dataset was denoised using the anatomical component based correction method (aCompCor) (Behzadi et al., 2007). Alternatively, the ME-uncleaned dataset was denoised using Automatic Removal of Motion Artifacts (ICA-AROMA, aggressive option) (Pruim et al., 2015).

**Figure 1.**
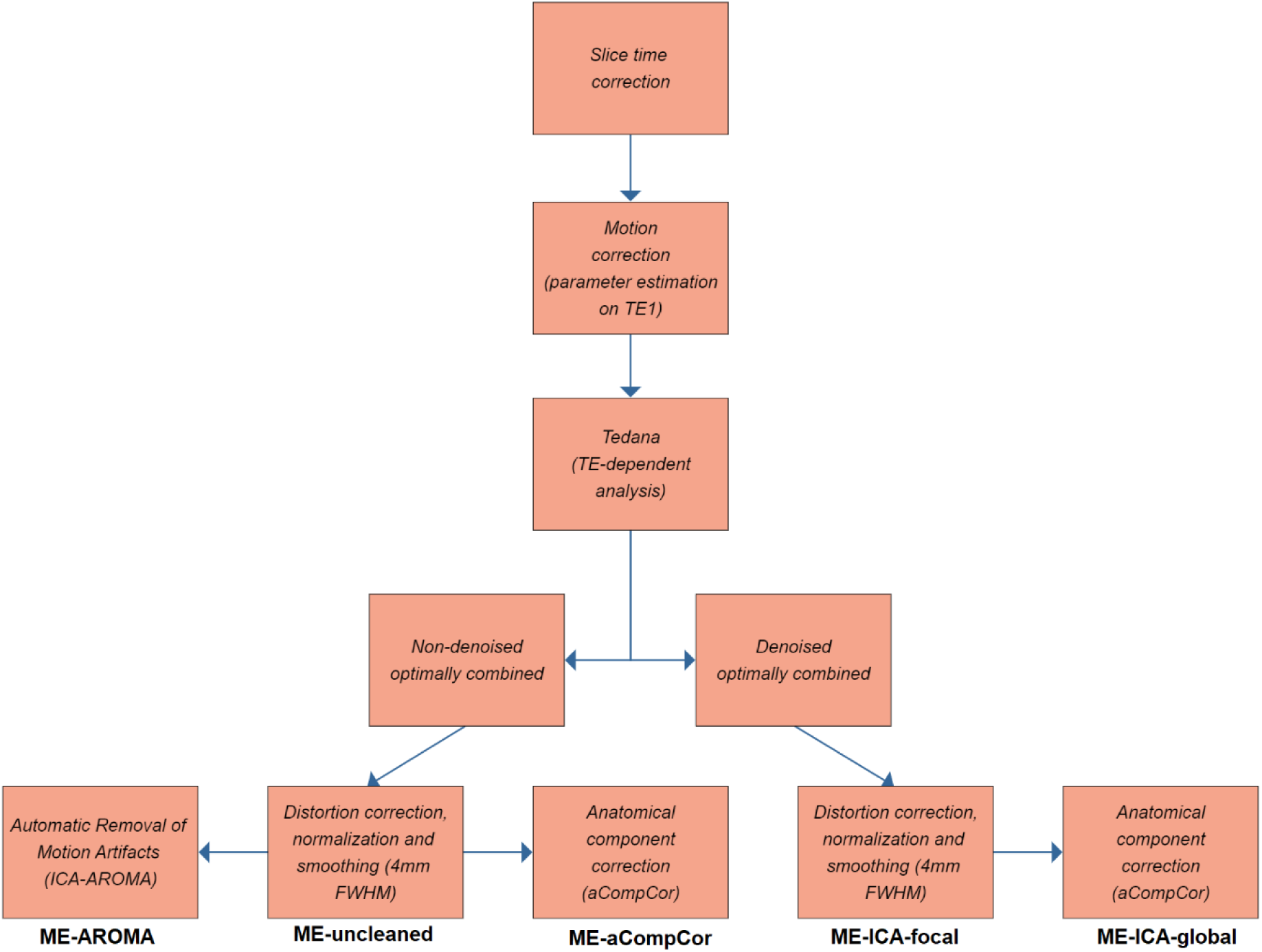
Flowchart demonstrating preprocessing pipeline, which branches after TE-dependent analysis (*tedana*). Intensity normalization and nuisance regressors were applied after *tedana* in order to prevent distortion of T2* estimations. Dataset names are shown at the bottom of the flowchart: these names are referred to throughout the manuscript.

In addition to the non-denoised dataset, *tedana* provided an optimally combined dataset that was denoised using ME-ICA. To this end, principal component analysis followed by the stabilized Kundu component selection decision tree was applied to the optimally combined data for dimensionality reduction (Kundu et al., 2013). Independent component analysis was then used to decompose the dimensionally reduced dataset. Component selection was performed to identify BOLD (TE-dependent), non-BOLD (TE-independent), and uncertain (low-variance) components using the Kundu decision tree (Kundu et al., 2013). The resulting denoised dataset was corrected for susceptibility-induced distortion, segmented, normalized to MNI space and finally smoothed using a 4mm Gaussian kernel (similarly to the ME-uncleaned data). This dataset was considered rid of spatially focal noise (Power et al., 2018), and hereafter referred to as ME-ICA-focal. Spatially diffuse noise was subsequently removed from the ME-ICA-focal dataset using aCompCor, and hereafter referred to as ME-ICA-global. Finally, a high pass filter of 0.008 Hz (125 seconds) was applied to all datasets.

### Data quality assessment

In order to assess data quality, various quality metrics were calculated and compared between denoising methods.

1. Temporal Signal-to-Noise Ratio (tSNR) was calculated for all datasets using the formula 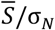 where 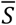 represents the mean fMRI signal and σN the standard deviation (SD) of the noise. The mean tSNR (and SD) was calculated across subjects for each cleaning method. This was performed first using a subject specific grey matter mask. Additionally, tSNR was calculated using a brainstem mask. Finally, tSNR ratio maps were created, where the group level tSNR map of each denoising method was divided by the tSNR map of the ME-uncleaned dataset. These tSNR ratio maps were used to identify which regions benefited the most of each respective data cleaning method.
2. Delta variation signal (DVARS), i.e. the frame-to-frame root mean square change in fMRI signal was calculated using the formula 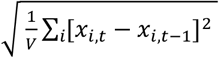 where *V* is the number of non-null voxels and *x* the fMRI signal in voxel *i* at time *t*, as per (Afyouni and Nichols, 2018). DVARS was used to identify and compare artefactual signal changes in the fMRI time series – likely originating from subject motion – that remained after each data cleaning method. A qualitative comparison between DVARS for each data cleaning method and subject motion was performed at the subject level using a framewise displacement calculation (Power et al., 2012).
3. Power spectral density was estimated using the multitaper method as available in MATLAB 2018a. Power spectra were calculated from fMRI time series at the subject level for each data cleaning method. Power spectra were subsequently normalized to the amplitude of the respective uncleaned series. Finally, power spectra were averaged for each data cleaning method. An adequate denoising method was expected to suppress non-neuronal frequencies (> 0.1 Hz (He et al., 2008)) while simultaneously preserving power spectral density amplitude in the low frequency range (< 0.1 Hz).
4. It has been demonstrated previously that the respiratory cycle correlates significantly with the global signal (Power et al., 2018). This has been reported to be due to a) changes in blood gasses and b) motion related to respiration. Adequate denoising methods should uncouple the global BOLD signal from respiration. Correlation between the variability in the global signal and the variability in the respiratory cycle was calculated as described by (Power et al., 2018). Signals from the respiratory belt recordings were z-scored, and the envelope of each signal was calculated using the built-in MATLAB function (‘peak’ method). Finally, the standard deviation of the respiratory envelope was calculated to capture variability in respiratory patterns.

## Results

### Temporal Signal-to-Noise Ratio (tSNR)

The tSNR values per data cleaning method are plotted in Figure 2 for whole-brain and brainstem separately (panel A and B, respectively). Each data cleaning method significantly improved tSNR, as shown in Table 1. The highest tSNR was achieved with the use of AROMA. Slightly lower tSNR values (as compared to whole-brain) were found when applying a brainstem mask (Figure 2B), although overall results were comparable, including the highest value for AROMA. Subsequently, we assessed the improvement in tSNR as compared to the uncleaned data using tSNR ratio maps (Figure 3). A relative uniform increase throughout the brain in tSNR was found with the use of aCompCor. For ME-ICA-focal, ME-ICA-global and ME-AROMA, the increase in tSNR was most apparent around the edges of the brain and across tissue boundaries (e.g. around the ventricles). When looking into the relative improvement in tSNR for whole-brain and brainstem separately, we found that for ME-ICA-focal and ME-ICA-global gain in tSNR was higher in the brainstem as compared to whole-brain (Table 2). ME-AROMA showed a higher gain in tSNR at the whole-brain level as compared to the brainstem. ME-aCompCor showed comparable gains in tSNR at the whole-brain and brainstem level.

**Figure 2.**
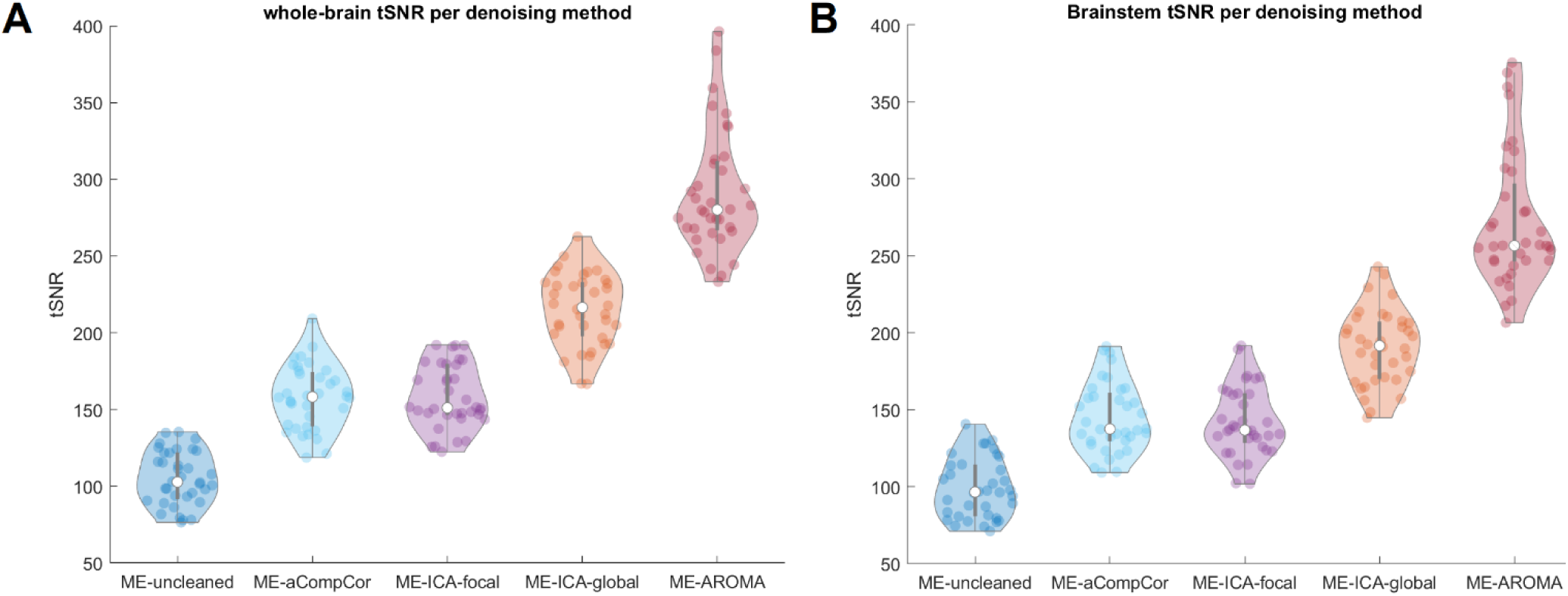
Violin plots of tSNR values per data cleaning method, panel A: tSNR at whole-brain level, panel B: tSNR at brainstem level.

**Table 1.** Comparison of tSNR subject means per data cleaning method (versus uncleaned data) using a paired-samples T-test.

|  |  |  |
| --- | --- | --- |
| ME-aCompCor > ME-uncleaned | t(35) = 19,90 | P < 0.001 |
| ME-ICA-focal > ME-uncleaned | t(35) = 23,93 | P < 0.001 |
| ME-ICA-global > ME-uncleaned | t(35) = 40,39 | P < 0.001 |
| ME-AROMA > ME-uncleaned | t(35) = 34,60 | P < 0.001 |
| ME-ICA-focal > ME-aCompCor | t(35) = 0,03 | P = 0.97 |
| ME-ICA-global > ME-ICA-focal | t(35) = 24,50 | P < 0.001 |
| ME-AROMA > ME-ICA-global | t(35) = 15,53 | P < 0.001 |

**Figure 3.**
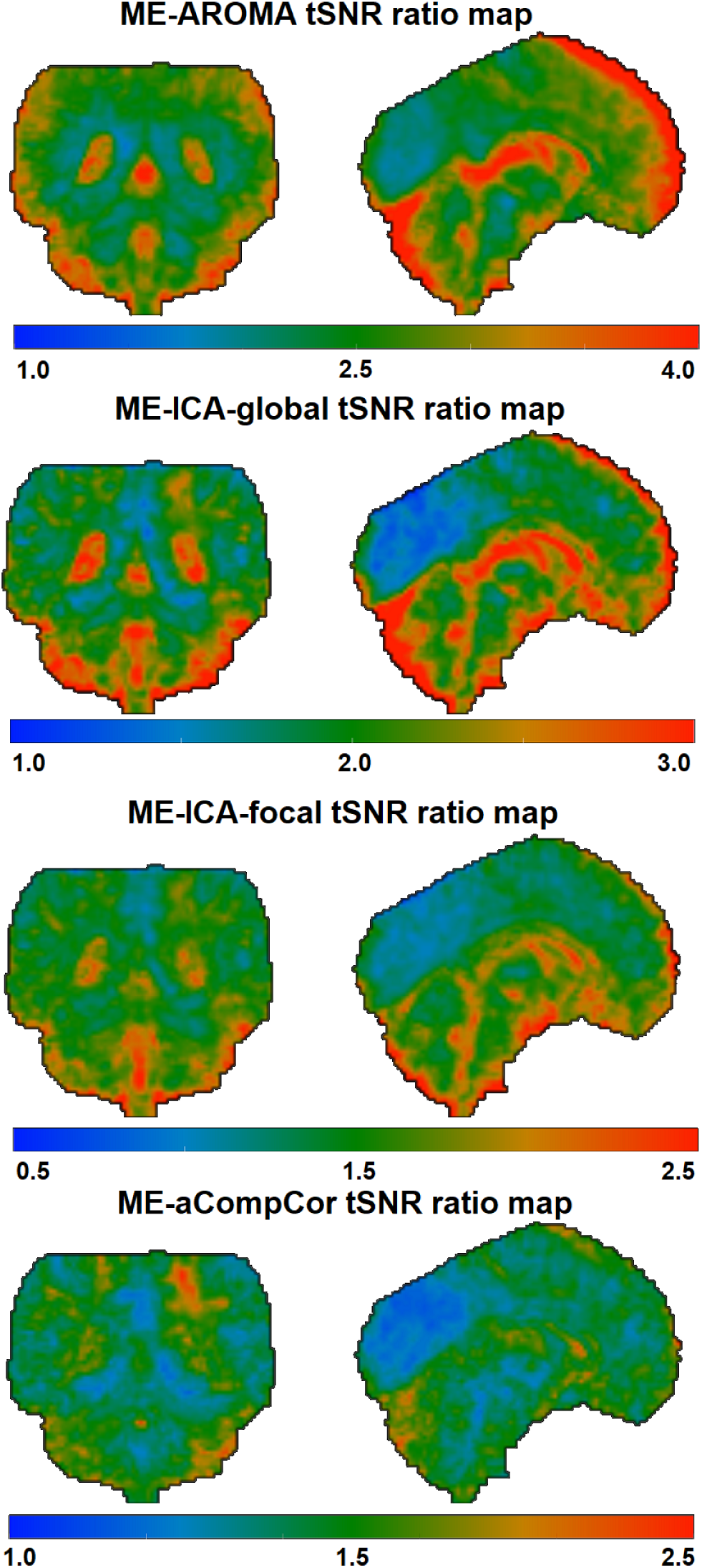
tSNR ratio maps: tSNR maps for each data cleaning method are divided by the tSNR map of the ME-uncleaned dataset. Color scale ranges from blue to green to red (low – high increase in tSNR, respectively).

**Table 2.** Relative improvement in tSNR at the whole-brain and brainstem level, as compared to ME-uncleaned (uncleaned data)

|  |  |  |
| --- | --- | --- |
|  | Whole-brain ratio | Brainstem ratio |
| ME-aCompCor | 1.43 | 1.46 |
| ME-ICA-focal | 1.57 | 1.74 |
| ME-ICA-global | 2.13 | 2.23 |
| ME-AROMA | 2.90 | 2.53 |

### Delta variation signal (DVARS)

The variability (standard deviation) of the DVARS per data cleaning method is displayed in Figure 4. Each data cleaning method significantly improved DVARS, as shown in Table 3. The lowest variability in DVARS was found when using AROMA.

**Figure 4.**
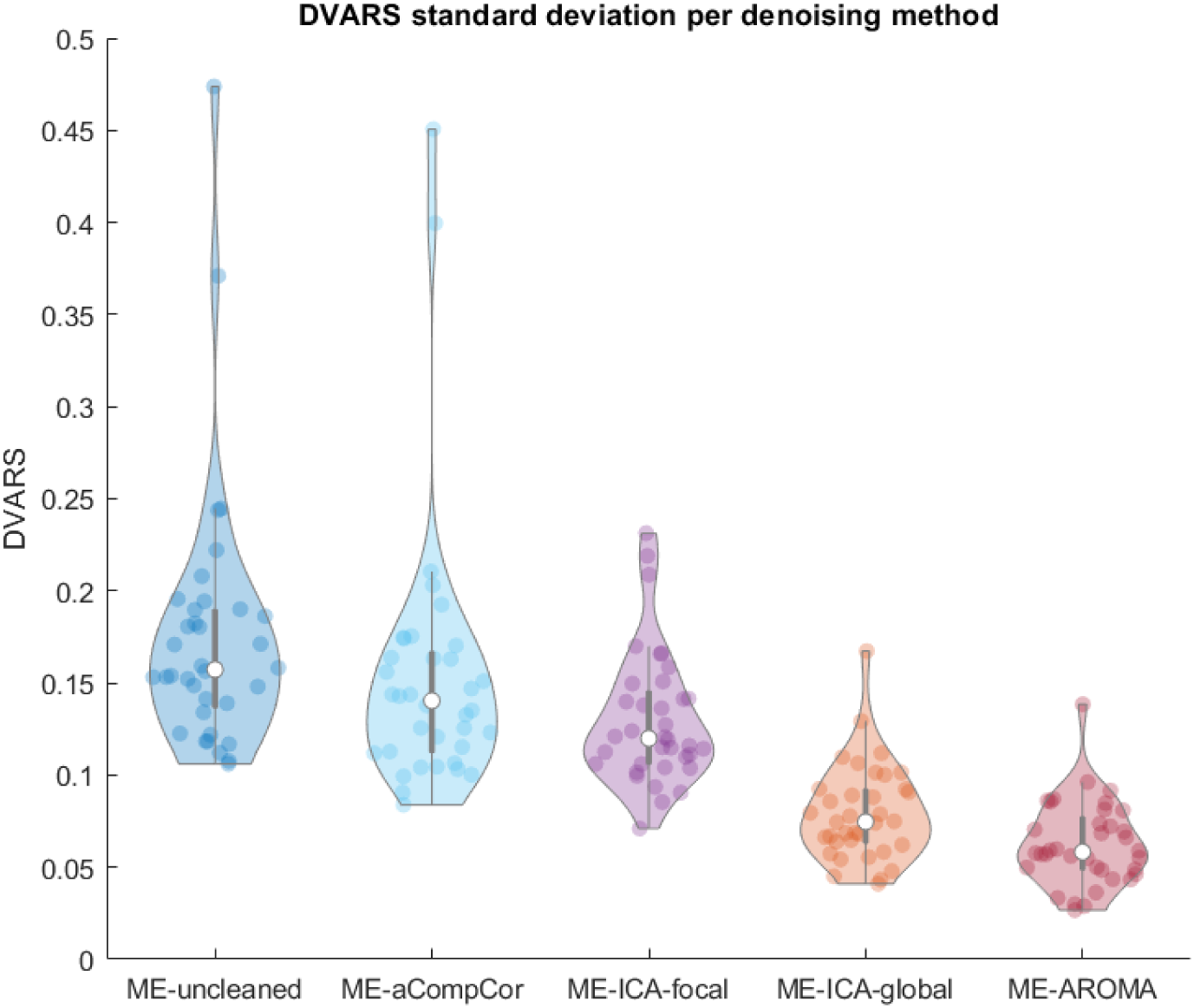
Violin plot of DVARS standard deviation values per data cleaning method.

**Table 3.** Comparison of DVARS subject means per data cleaning method using a paired-samples T-test.

|  |  |  |
| --- | --- | --- |
| ME-aCompCor > ME-uncleaned | t(35) = 7,62 | P < 0.001 |
| ME-ICA-focal > ME-uncleaned | t(35) = 23,32 | P < 0.001 |
| ME-ICA-global > ME-uncleaned | t(35) = 26,26 | P < 0.001 |
| ME-AROMA > ME-uncleaned | t(35) = 26,75 | P < 0.001 |
| ME-ICA-focal > ME-aCompCor | t(35) = 15,18 | P < 0.001 |
| ME-ICA-global > ME-ICA-focal | t(35) = 10,31 | P < 0.001 |
| ME-AROMA > ME-ICA-global | t(35) = 8,57 | P < 0.001 |

In order to make a qualitative comparison of artefactual signal changes remaining after each data cleaning method, we plotted DVARS with framewise displacement (displacement between neighboring voxels) for two individual subjects: one subject demonstrating low movement during scanning (mean framewise displacement of 0.06 mm) and one subject demonstrating high movement (mean framewise displacement of 0.19 mm). DVARS was plotted as timeseries for each data cleaning method; peaks in signal change can be compared to peaks in framewise displacement (Figure 5).

**Figure 5.**
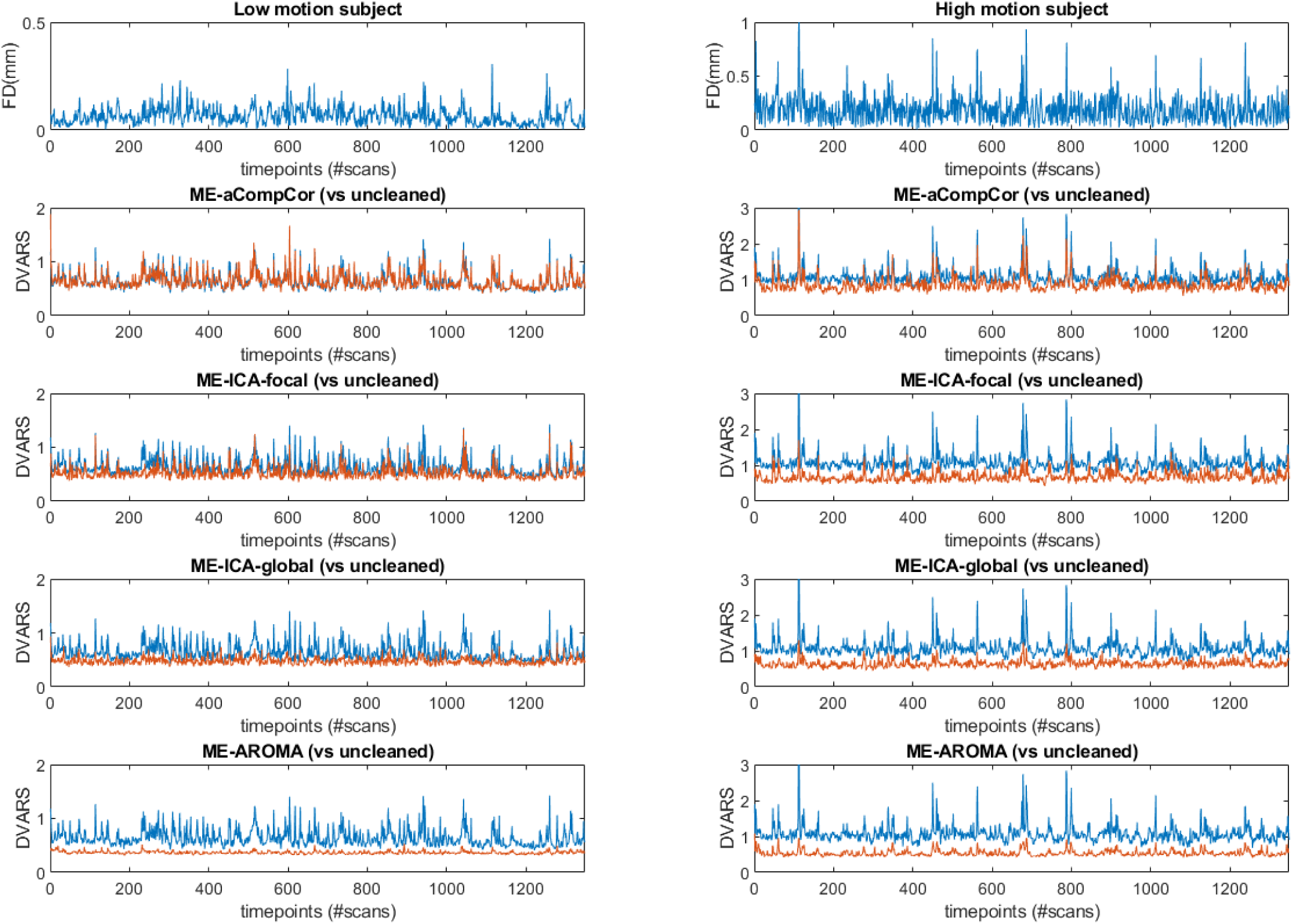
Upper two panels display framewise displacement (FD): panels on the left correspond to a subject demonstrating low movement (FD = 0.06 mm), panels on the right correspond to a subject demonstrating high movement (FD = 0.19 mm). DVARS is plotted for each data cleaning method (red) against DVARS for the uncleaned data (blue).

### Power Spectral Density

Power spectra averaged per data cleaning method are displayed in Figure 6. Data cleaning with ME-ICA only (ME-ICA-focal) had little impact on the power spectrum both in the low and high frequency range. Data cleaning with aCompCor had a marked impact on the power amplitude at frequencies > 0.1 Hz, whilst maintaining the peak in the power spectrum at frequencies around 0.02 Hz (within the BOLD frequency range).

**Figure 6.**
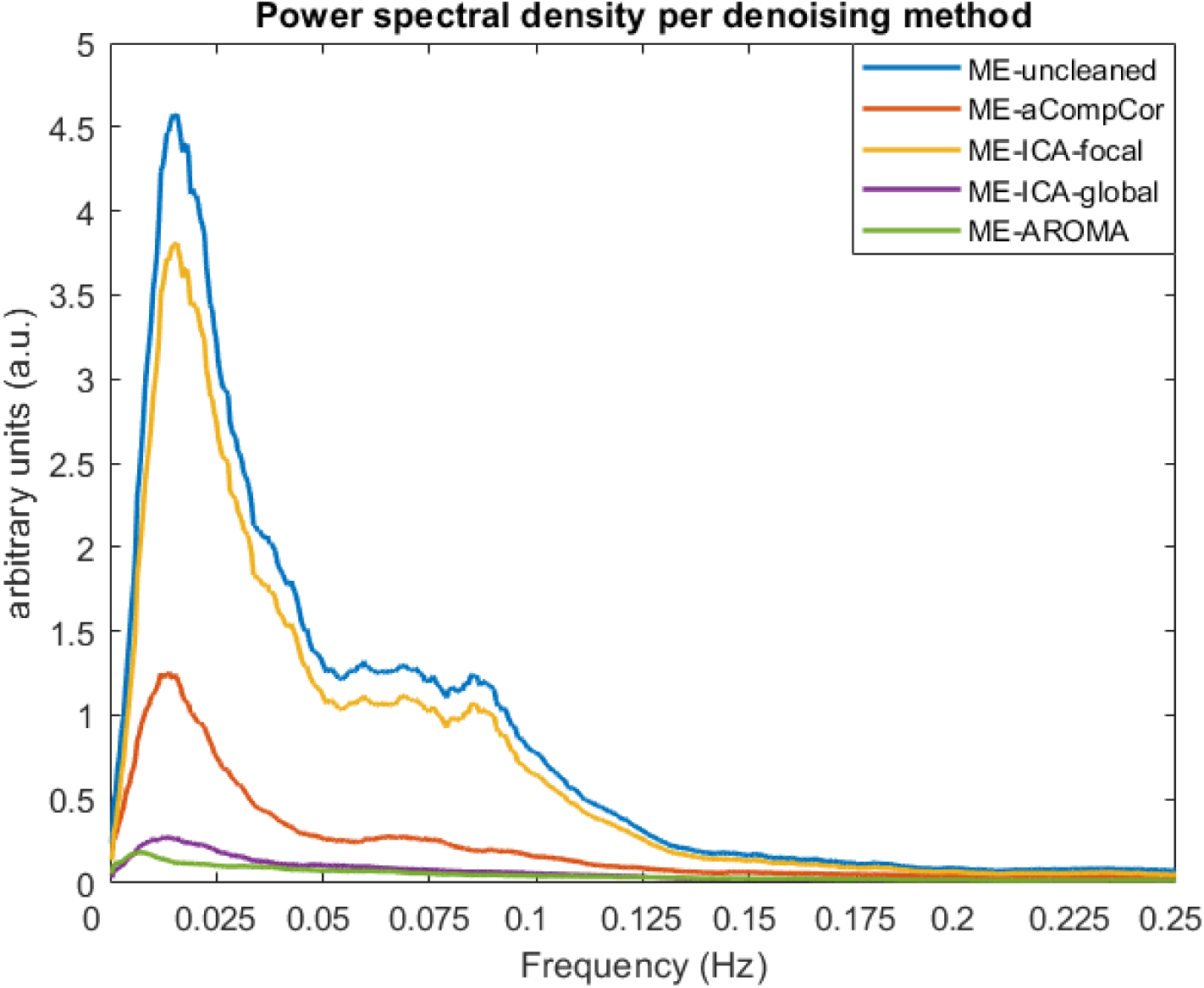
Power spectra averaged per data cleaning method.

**Figure 7.**
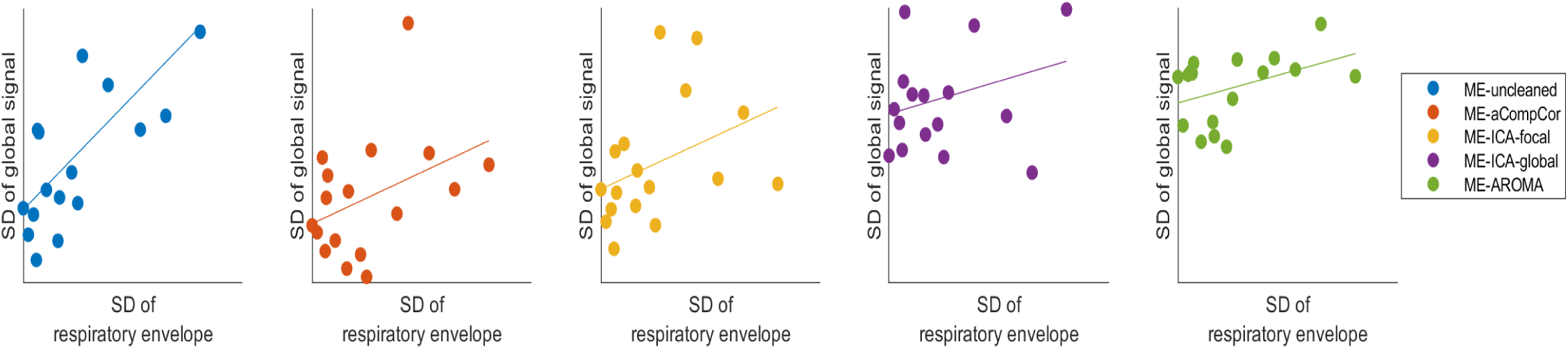
Correlation between respiration and global signal. Figure shows scatterplots and least-square lines with on the y-axis the variability (standard deviation) of the global signal of each dataset and on the x-axis the variability of the respiratory cycle, calculated as the standard deviation of the envelope of the z-scored respiratory belt signal.

Frequencies above > 0.1 Hz were further suppressed when combining ME-ICA with aCompCor (ME-ICA-global), though at the cost of spectral density in the BOLD frequency range. With the use of AROMA, the power spectrum in both the low and high frequency range was heavily suppressed.

### Correlation between breathing and global signal

The correlation between the variability of the global signal and variability of the respiratory envelope was calculated for each dataset. A strong correlation was found for the uncleaned dataset (r = 0.70, p = 0.002). No significant correlation was found between global signal and respiratory envelope for any of the cleaned datasets: ME-aCompCor (r = 0.40, p = 0.111), ME-ICA-focal (r = 0.39, p = 0.127), ME-ICA-global (r = 0.28, p = 0.271), ME-AROMA (r = 0.42, p = 0.095).

## Discussion

The current study is the first to investigate the efficacy of data cleaning methods of multi-echo fMRI data acquired at 7T. Given the absence of a so-called ground truth when using BOLD data one can only examine quality measures related to the data itself. In the current study we have used several measures that are largely unrelated to each other, with each measure highlighting different aspects of the data. We here demonstrate significant tSNR improvements with each of the used data cleaning methods. tSNR improvements showed a similar pattern for whole-brain and brainstem data only. AROMA resulted in the highest tSNR, followed by ME-ICA combined with aCompCor (ME-ICA-global). Notably, the tSNR ratio maps showed that for both the ME-AROMA and ME-ICA-global dataset, tSNR improvements were strongest around the edges of the brain and brainstem. This is in line with the previous finding that motion artefacts are most profound across the edges of the brain (Patriat et al., 2015; Satterthwaite et al., 2013). Moreover, differences in proton density across different tissues, for example around the ventricles, similarly result in increased noise (Caballero-Gaudes and Reynolds, 2017). Denoising is therefore likely to have a stronger effect on these regions.

When comparing variability in DVARS, we found the lowest values for AROMA. In a qualitative analysis, it appears that denoising has a stronger effect on DVARS for subjects with higher degrees of motion, likely due to higher baseline signal variation. It should be noted that strong peaks related to motion remain visible in the DVARS timecourse even after applying ME-ICA or AROMA. One might therefore consider using additional steps such as scrubbing in order to censor remaining artefactual scanning volumes, which is how we preprocessed the data for our primary analysis, as discussed elsewhere (Beckers et al., 2021).

We subsequently compared power spectral density of each dataset. Whilst tSNR and DVARS were superior for ME-AROMA and ME-ICA-global, the power spectrum of these datasets indicated a strong reduction of both high (non-neuronal) and low (neuronal) frequencies, with the spectrum being most suppressed after applying AROMA. It is therefore possible that, although AROMA is highly effective in removing noise, the approach is too aggressive in the sense that it also removes signal of interest. It should be noted that AROMA can be performed with more lenient settings, as incorporated in the software itself (“non-aggressive” option). Due to the data size and number of comparisons, we have not used both options of AROMA here; hence, we cannot draw conclusions on data quality after running AROMA with the non-aggressive option. It is possible that the non-aggressive option would retain more signal of interest, but this would likely also result in less effective removal of noise. This was also demonstrated in a study comparing data cleaning methods applied to single-echo data acquired at 1.5T, including both options of AROMA (Dipasquale et al., 2017). This study furthermore applied the aggressive option of AROMA to multi-echo data, comparing it to ME-ICA. Given that resting state fMRI data was used, the authors compared connectivity maps after each data cleaning method. The authors stated that ME-ICA more effectively preserved functional connectivity (i.e. signal of interest) as compared to AROMA.

We finally looked at the ability of each data cleaning method to uncouple the global signal and respiratory cycle, as it has previously been demonstrated that respiration strongly affects the BOLD signal (Power et al., 2018). We indeed found a strong correlation when analyzing the uncleaned dataset. After each data cleaning method, the correlation between the global signal and respiratory cycle was no longer significant. This contrasts the previous finding of Power et al, where the correlation between global signal and respiration remained after applying ME-ICA without additional global signal regression or aCompCor. This discrepancy could be both data and software related. First, the study by Power and colleagues was performed using 3T data. The proportion of physiological noise increases substantially with higher field strengths (Triantafyllou et al., 2005), which could affect the relationship between the global signal and respiratory cycle. Moreover, the software for ME-ICA has evolved over time. The first versions were developed to function using AFNI modules (Kundu et al., 2012), whereas the version used here (*tedana*) is a standalone python-based program.

## Conclusions

AROMA and ME-ICA combined with aCompCor encompass highly effective denoising options for multi-echo 7T fMRI data, as compared to the commonly used aCompCor. Combining ME-ICA and aCompCor appears to have an additive effect based on the quality metrics used here, making it superior to ME-ICA alone. When using AROMA with the aggressive denoising option, it is possible that a portion of signal of interest is falsely classified as noise. When using AROMA, empirical testing of both the aggressive and non-aggressive option per dataset might therefore be the best choice.

## Acknowledgements

We thank the Scannexus support team for their assistance during data acquisition. This study was in part funded by grants from the Brains Unlimited Pioneer Fund and ‘Stichting Sint Annadal’ Foundation.

## Disclosures

DK has received grants from Will Pharma, Allergan, Grunenthal, ZonMw, MLDS and UEG, outside of submitted work. The authors state no conflicting interests.

## Author Contributions

**Abraham B Beckers:** Conceptualization; Data curation; Formal analysis; Investigation; Methodology; Project administration; Software; Validation; Visualization; Writing - original draft. **Benedikt A Poser:** Conceptualization; Methodology; Software; Writing - review & editing. **Daniel Keszthelyi**: Conceptualization; Funding acquisition; Resources; Supervision; Writing - review & editing.

## Data Availability Statement

All custom MATLAB code and metadata used for the analyses is available in the following GitHub repository: https://github.com/BramBeckers/7TMulti-echo. The raw datasets analyzed during the current study are available from the corresponding author on reasonable request.

## Notes

### Competing Interest Statement

The authors have declared no competing interest.

## References

Afyouni, S., Nichols, T.E., 2018. Insight and inference for DVARS. Neuroimage 172, 291–312.

Andersson, J.L., Skare, S., Ashburner, J., 2003. How to correct susceptibility distortions in spin-echo echo-planar images: application to diffusion tensor imaging. Neuroimage 20, 870–888.

Beckers, A.B., van Oudenhove, L., Weerts, Z., Jacobs, H.I.L., Priovoulos, N., Poser, B.A., Ivanov, D., Gholamrezaei, A., Aziz, Q., Elsenbruch, S., Masclee, A.A.M., Keszthelyi, D., 2021. Evidence for engagement of the nucleus of the solitary tract in processing intestinal chemonociceptive input irrespective of conscious pain response in healthy humans. Pain.

Behzadi, Y., Restom, K., Liau, J., Liu, T.T., 2007. A component based noise correction method (CompCor) for BOLD and perfusion based fMRI. Neuroimage 37, 90–101.

Brooks, J.C., Faull, O.K., Pattinson, K.T., Jenkinson, M., 2013. Physiological noise in brainstem FMRI. Front Hum Neurosci 7, 623.

Caballero-Gaudes, C., Reynolds, R.C., 2017. Methods for cleaning the BOLD fMRI signal. Neuroimage 154, 128–149.

Dipasquale, O., Sethi, A., Lagana, M.M., Baglio, F., Baselli, G., Kundu, P., Harrison, N.A., Cercignani, M., 2017. Comparing resting state fMRI de-noising approaches using multi-and single-echo acquisitions. PLoS One 12, e0173289.

Glover, G.H., Li, T.Q., Ress, D., 2000. Image-based method for retrospective correction of physiological motion effects in fMRI: RETROICOR. Magn Reson Med 44, 162–167.

He, B.J., Snyder, A.Z., Zempel, J.M., Smyth, M.D., Raichle, M.E., 2008. Electrophysiological correlates of the brain’s intrinsic large-scale functional architecture. Proc Natl Acad Sci U S A 105, 16039–16044.

Kong, Y., Jenkinson, M., Andersson, J., Tracey, I., Brooks, J.C., 2012. Assessment of physiological noise modelling methods for functional imaging of the spinal cord. Neuroimage 60, 1538–1549.

Kundu, P., Inati, S.J., Evans, J.W., Luh, W.M., Bandettini, P.A., 2012. Differentiating BOLD and non-BOLD signals in fMRI time series using multi-echo EPI. Neuroimage 60, 1759–1770.

Kundu, P., Voon, V., Balchandani, P., Lombardo, M.V., Poser, B.A., Bandettini, P.A., 2017. Multi-echo fMRI: A review of applications in fMRI denoising and analysis of BOLD signals. Neuroimage 154, 59–80.

Kwong, K.K., Belliveau, J.W., Chesler, D.A., Goldberg, I.E., Weisskoff, R.M., Poncelet, B.P., Kennedy, D.N., Hoppel, B.E., Cohen, M.S., Turner, R., et al., 1992. Dynamic magnetic resonance imaging of human brain activity during primary sensory stimulation. Proc Natl Acad Sci U S A 89, 5675–5679.

Liu, T.T., 2016. Noise contributions to the fMRI signal: An overview. Neuroimage 143, 141–151.

Napadow, V., Sclocco, R., Henderson, L.A., 2019. Brainstem neuroimaging of nociception and pain circuitries. Pain Rep 4, e745.

Patriat, R., Molloy, E.K., Birn, R.M., 2015. Using Edge Voxel Information to Improve Motion Regression for rs-fMRI Connectivity Studies. Brain Connect 5, 582–595.

Poser, B.A., Norris, D.G., 2009. Investigating the benefits of multi-echo EPI for fMRI at 7 T. Neuroimage 45, 1162–1172.

Posse, S., Wiese, S., Gembris, D., Mathiak, K., Kessler, C., Grosse-Ruyken, M.L., Elghahwagi, B., Richards, T., Dager, S.R., Kiselev, V.G., 1999. Enhancement of BOLD-contrast sensitivity by single-shot multi-echo functional MR imaging. Magn Reson Med 42, 87–97.

Power, J.D., Barnes, K.A., Snyder, A.Z., Schlaggar, B.L., Petersen, S.E., 2012. Spurious but systematic correlations in functional connectivity MRI networks arise from subject motion. Neuroimage 59, 2142–2154.

Power, J.D., Plitt, M., Gotts, S.J., Kundu, P., Voon, V., Bandettini, P.A., Martin, A., 2018. Ridding fMRI data of motion-related influences: Removal of signals with distinct spatial and physical bases in multiecho data. Proc Natl Acad Sci U S A 115, E2105–E2114.

Pruim, R.H.R., Mennes, M., van Rooij, D., Llera, A., Buitelaar, J.K., Beckmann, C.F., 2015. ICA-AROMA: A robust ICA-based strategy for removing motion artifacts from fMRI data. Neuroimage 112, 267–277.

Puckett, A.M., Bollmann, S., Poser, B.A., Palmer, J., Barth, M., Cunnington, R., 2018. Using multi-echo simultaneous multi-slice (SMS) EPI to improve functional MRI of the subcortical nuclei of the basal ganglia at ultra-high field (7T). Neuroimage 172, 886–895.

Satterthwaite, T.D., Elliott, M.A., Gerraty, R.T., Ruparel, K., Loughead, J., Calkins, M.E., Eickhoff, S.B., Hakonarson, H., Gur, R.C., Gur, R.E., Wolf, D.H., 2013. An improved framework for confound regression and filtering for control of motion artifact in the preprocessing of resting-state functional connectivity data. Neuroimage 64, 240–256.

Speck, O., Hennig, J., 1998. Functional imaging by I0-and T2*-parameter mapping using multi-image EPI. Magn Reson Med 40, 243–248.

Triantafyllou, C., Hoge, R.D., Krueger, G., Wiggins, C.J., Potthast, A., Wiggins, G.C., Wald, L.L., 2005. Comparison of physiological noise at 1.5 T, 3 T and 7 T and optimization of fMRI acquisition parameters. Neuroimage 26, 243–250.

Whitfield-Gabrieli, S., Nieto-Castanon, A., 2012. Conn: a functional connectivity toolbox for correlated and anticorrelated brain networks. Brain Connect 2, 125–141.

